# MuLAN: Mutation-driven Light Attention Networks for investigating protein-protein interactions from sequences

**DOI:** 10.1101/2024.08.24.609515

**Authors:** Gianluca Lombardi, Alessandra Carbone

## Abstract

Understanding how proteins interact and how mutations affect these interactions is crucial for unraveling the complexities of biological systems and their evolution. Mutations can significantly alter protein behavior, impacting stability, interactions, and activity, thereby affecting cellular functions and influencing disease development and treatment effectiveness. Experimental methods for examining protein interactions are often slow and costly, highlighting the need for efficient computational strategies. We present MuLAN, a groundbreaking deep learning method that leverages light attention networks and the power of pre-trained protein language models to infer protein interactions, predict binding affinity changes, and reconstruct mutational landscapes for proteins involved in binary interactions, starting from mutational changes and directly using sequence data only. Unlike previous methods that depend heavily on structural information, MuLAN’s sequence-based approach offers faster and more accessible predictions. This innovation allows for variations in predictions based on specific partners, opening new possibilities for understanding protein behavior through their sequences. The potential implications for disease research and drug development mark a significant step forward in the computational analysis of protein interactions.

## Introduction

Proteins are vital to cell function, and understanding their behaviour during interactions within complex systems is key to deciphering the biological mechanisms of life. This task becomes particularly daunting when considering how mutations affect protein interactions and their binding strength. Such mutations can impact protein behavior in several ways, including altering stability, affecting functional activity, and modifying protein-protein interactions (PPIs). These changes are typically triggered by physical alterations at the interaction sites, such as changes in hydrophobic areas, electrostatic interactions, or hydrogen bonds, leading to variations in the binding free energy, Δ*G*, between mutated and wild-type protein complexes. The assessment of this variance, or “binding affinity change” ΔΔ*G*, is crucial for understanding disease predisposition and the effectiveness of drugs (Kastritis and Bonvin, 2013).

Traditionally, experimental techniques have been employed to examine protein complex binding affinities, but these methods are expensive and time-consuming (Geng *et al*., 2019a). This has spurred the development of computational strategies to bypass these drawbacks. Over the past decade, numerous computational approaches considering three dimensional structure information have been introduced to assess binding energy differences. They include molecular dynamics simulations reaching very accurate estimations, but requiring extensive computational resources which limit their large-scale application (Siebenmorgen and Zacharias, 2020). To address these challenges, empirical energy functions like FoldX and Rosetta have been developed. Despite their faster predictions, these models suffer from limited conformational sampling, leading to reduced accuracy. Recently, advancements in deep learning based on structural data have shown promise, though their effectiveness heavily relies on the availability of extensive training datasets reporting experimental data for known complex structures. The creation of the SKEMPI v2.0 dataset (Jankauskait *et al*., 2018) has significantly enhanced the precision and efficiency of methods like iSEE (Geng *et al*., 2019b), mCSM-PPI2 (Rodrigues *et al*., 2019), BindProfX (Xiong *et al*., 2017), MuPIPR (Zhou *et al*., 2020), MutaBind2 (Zhang *et al*., 2020), GeoPPI (Liu *et al*., 2021), and DLA-Mutations (Mohseni Behbahani *et al*., 2023). All these methods incorporate geometrical and physico-chemical information on the complexes, and in some cases, information on the changes in binding mechanisms, involving atomic properties, which are known to directly contribute to the binding affinity. A fundamental bottleneck on the performance of all computational methods so far is the availability of high quality structural complexes for both wild-type and mutants upon which they rely. The lack of suitable complex structures affects the large scale application of the methods (Xiong *et al*., 2022).

The advent of protein language models and deep learning architectures, such as transformers (Vaswani *et al*., 2017), presents a novel challenge: understand protein interactions from sequences. Achieving this would usher in a new era, offering fresh insights into protein behavior based solely on their sequences and providing faster computational predictions. We hypothesize that protein mutational changes are key to inferring protein interaction sites and reconstructing mutational landscapes for proteins involved in binary interactions, directly using sequence data. To address this, we introduce a deep learning approach, MuLAN, based on light attention layers that combine contextualized representations from pre-trained protein language models to encode protein complexes and predict binding affinity changes due to single-point and multiple-point mutations. As anticipated, MuLAN’s predictions will facilitate a detailed understanding of protein interactions (Figure 1).

**Figure 1:**
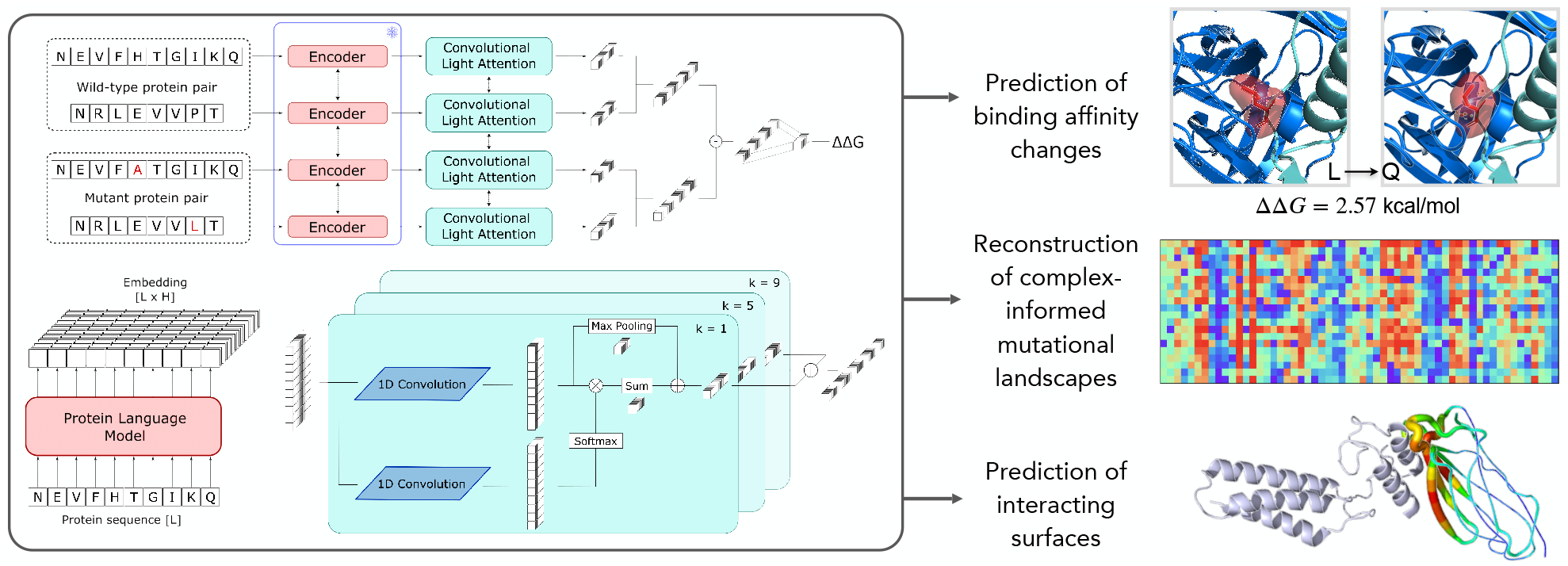
Representation of MuLAN model and its tasks. The model leverages a Protein Language Model encoder to extract features from wild-type and mutated sequences within a complex, and processes them through a convolutional light attention block, that produces interpretable representations for the prediction. The MuLAN architecture (left) is employed for three main goals (right): 1. to predict variations in binding affinity (top); 2. to reconstruct entire complex-informed mutational landscapes (center); 3. to identify potential interacting surfaces from the representations learned by the attention mechanism (bottom).

## Results

We introduce a deep learning architecture, MuLAN, which takes four sequences as input: two representing a wild-type binary complex and two representing the mutated sequences within the complex. MuLAN outputs a score evaluating the variations in binding affinity (ΔΔ*G*) caused by single or few-point mutations, potentially occurring in both sequences. MuLAN is central to our approach for understanding protein interactions from sequences. It achieves three main goals: predicting binding affinity changes, identifying residues at the protein interface within the complex, and reconstructing complex-informed mutational landscapes. This last goal marks significant progress over previous methods, as it uniquely accounts for the dependency of the mutational landscape on the interacting protein partner. The role of interaction comes to the forefront of our protein analysis, highlighting the crucial influence of specific protein partners on mutational outcomes. This emphasis on interaction is a critical aspect that has been ignored by current computational methods, making MuLAN approach innovative and impactful.

Figure 1 illustrates the design of the MuLAN architecture (see Methods). The design is guided by the principle that predicting properties related to protein complexes solely from sequences, while maintaining plausible biological interpretations, requires a model that processes input sequences and integrates their representations without losing information about individual residues. MuLAN embodies this intuition by learning a weighted average of residue embeddings extracted from pre-trained language models for both wildtype and mutated protein sequences to predict binding affinity changes caused by mutations. To manage the high dimensionality of the embeddings, we first apply parallel convolutional filters to learn a lower-dimensional representation, effectively combining information from neighboring residues. Light attention is achieved by applying softmax normalization to the convolution output over the sequence length, with computational complexity scaling linearly with sequence length. It is important to note that long-range correlations, which cannot be captured by this approach, are already encoded in the sequence embeddings. For ΔΔ*G* prediction, we use a multi-layer perceptron with linear activation on the difference of embeddings for mutated and wild-type complexes. In addition to the baseline MuLAN model, we trained a second version called *i*MuLAN. This enhanced model incorporates iGEMME scores (Tekpinar *et al*., 2024) as auxiliary features, representing the sensitivity of mutations to be deleterious. These scores were concatenated to the input of the final linear layer in MuLAN. Since iGEMME scores pertain to single-point mutations, we evaluated iMuLAN exclusively for this scenario.

### MuLAN predicts binding affinity changes in binary complexes

Binding affinity changes can be predicted from the sequences of both wild-type and mutant proteins. A comprehensive assessment of MuLAN’s performance was conducted through a series of experiments involving thousands of mutations and hundreds of complexes. We used benchmark datasets for both single and multiple mutations. These experiments utilized datasets derived from SKEMPI v2.0, including S1102, S1131, S2003, and S4169 for single-point mutations in binary complexes, and S1400 for multiple-point mutations. A comprehensive description of these datasets can be found in Methods and Table S1. We used Pearson’s correlation coefficient (PCC) and root mean squared error (RMSE) to compare the experimental values with those predicted by our model, following standard regression analysis practices. We conducted crossvalidation experiments using two distinct setups for splitting the training and testing data. In the first setup, a mutation-based split was used, where the training and testing data were divided according to individual mutations, allowing the model to encounter the same complexes in both phases but with different mutations. In the second setup, a complex-based split was employed, ensuring that the complexes used in training were entirely different from those in testing. This complex-based approach is more rigorous and provides a better assessment of the model’s generalizability to new, unseen sequences.

#### Mutation-based splitting

Both the baseline MuLAN model (Methods) and the *i*MuLAN version were evaluated using two different protein language models (pLMs) for extracting embeddings: ESM2–3B (Lin *et al*., 2023) and Ankh–large (Elnaggar *et al*., 2023). Following previous studies, we implemented a 10-fold cross-validation for datasets containing single-point mutations, and a 5-fold cross-validation for the S1400 dataset. Results are detailed in Table 1. The Ankh–large model outperformed the baseline across all datasets on both metrics, except for the PCC in the S4169 dataset. Notably, the embedding dimensions for these models are 2560 and 1536, respectively, indicating that embedding size is not the determining factor for the model’s predictive capability. Furthermore, integrating *i*GEMME scores consistently enhanced the performance metrics for both models, particularly in the S1102 and S1131 datasets.

**Table 1:** MuLAN and *i*MuLAN performance. RMSE is measured in kcal/mol.

| MODEL | S1102 |  | S1131 |  | S2003 |  | S4169 |  | S1400 |  |
| --- | --- | --- | --- | --- | --- | --- | --- | --- | --- | --- |
|  | PCC | RMSE | PCC | RMSE | PCC | RMSE | PCC | RMSE | PCC | RMSE |
| MUTATION-BASED SPLITTING |  |  |  |  |  |  |  |  |  |  |
| MuLAN-ESM2-3B | 0.854 | 1.223 | 0.849 | 1.255 | 0.781 | 1.078 | **0.739** | 1.216 | 0.904 | 1.187 |
| MuLAN-Ankh-large | **0.868** | **1.185** | 0.866 | 1.211 | 0.774 | 1.068 | **0.739** | 1.152 | **0.907** | **1.172** |
| *i*MuLAN-ESM2-3B | 0.864 | 1.193 | 0.858 | 1.248 | 0.779 | 1.086 | 0.725 | 1.152 | - | - |
| *i*MuLAN-Ankh-large | 0.867 | **1.185** | **0.871** | **1.196** | **0.792** | **1.059** | **0.739** | **1.134** | - | - |
| COMPLEX-BASED SPLITTING |  |  |  |  |  |  |  |  |  |  |
| *i*MuLAN-ESM2-3B | 0.698 | **1.469** | **0.733** | **1.469** | **0.685** | **1.320** | 0.564 | **1.357** | - | - |
| *i*MuLAN-Ankh-large | **0.714** | 1.480 | 0.731 | 1.495 | 0.618 | 1.417 | **0.568** | 1.419 | - | - |

MuLAN, which considers only the wild-type and mutated protein sequences as inputs, can easily handle multiple-point mutations — a capability often lacking in structure-based models. Notably, for the S1400 dataset, we achieved state-of-the-art results in terms of PCC and RMSE (Table 2). In the case of single-point mutations, our results were on par with other structure-based models for the S1102 and S1131 datasets. However, we observed lower PCC values for the S2003 and S4169 datasets, which are part of SKEMPI’s second release. We hypothesised that these lower scores were due to the embeddings from pLMs not capturing enough information about the properties and structures of some proteins in these datasets. To validate this hypothesis, we used ESMFold (Lin *et al*., 2023) to predict the structures of wild type proteins within the studied datasets and assessed model confidence using pTM and average pLDDT scores. We found that the average pTM scores were 0.83 for the S1102, S1131, and S1400 datasets, but lower for S4169 (0.74) and S2003 (0.73). Similarly, the average pLDDT scores were higher (0.78) for S1102, S1131, and S1400, and lower for S4169 (0.74) and S2003 (0.75). These lower scores for the S2003 and S4169 datasets suggest that the predicted structures were less reliable, potentially explaining why sequence information alone was insufficient for accurately predicting protein properties in these cases. This was particularly true for proteins in the antibody-antigen classes, where we observed a consistent drop in performance.

**Table 2:** MuLAN and *i*MuLAN performance on benchmark datasets from SKEMPI. RMSE is measured in kcal/mol. Missing values (“-”) are not reported by the corresponding analyses.

| MODEL | PCC | RMSE | MODEL | PCC | RMSE |
| --- | --- | --- | --- | --- | --- |
| S1400 |  |  | S2003 - mutation based |  |  |
| MuPIPR | 0.88 | 1.32 | DLA-Mutation | 0.81 | - |
| MuLAN | **0.91** | **1.17** | MuLAN | 0.79 | **1.06** |
| S1102 |  |  | S2003 - complex based |  |  |
| MuPIPR | 0.85 | 1.22 | DLA-Mutation | **0.735** | **1.23** |
| iSEE | 0.80 | 1.41 | DLA-Mutation* | 0.686 | 1.31 |
| MuLAN | **0.87** | **1.19** | MuLAN | 0.685 | 1.32 |
| S1131 |  |  | S4169 |  |  |
| TopNetTree | 0.85 | - | TopNetTree | **0.79** | - |
| Hom-ML-V2 | 0.857 | 1.28 | MpbPPI | **0.79** | - |
| MpbPPI | 0.865 | - | MuLAN | 0.74 | **1.13** |
| MuLAN | **0.87** | **1.20** |  |  |  |

#### Complex-based splitting

We conducted a 10-fold cross-validation on the single-point mutation datasets S1102, S1131, and S4169. For the S2003 dataset, we followed the complex-based split between training and testing sets as established in (Mohseni Behbahani *et al*., 2023). We excluded multiple-point mutations from this analysis and opted to evaluate using the *i*MuLAN version, which demonstrated superior metrics in mutation-based split scenarios. Although the evaluation metrics, particularly for the *i*MuLAN–Ankh–large model, tend to worsen in this setting, they still demonstrate the model’s strong generalization capabilities on unseen complexes (Table 1). Notably, the RMSE values remain below 1.5 kcal/mol across all datasets.

#### Comparison with other approaches

We compared ΔΔ*G* predictions of MuLAN with other recently developed approaches, previously evaluated on the same benchmark datasets extracted from SKEMPI. The majority of these methods rely on structural features for their predictions, or on a combination of sequence and structure information, while only MuPIPR (Zhou *et al*., 2020) is entirely sequence-based. On S1400, we achieve an higher PCC of 0.90 and a lower RMSE of 1.17 kcal/mol, with respect to MuPIPR reported values of 0.88 and 1.32 respectively, showing an improvement in predictions for multiple-point mutations, where other structure-based approaches typically struggle. On single-point mutations from S1102, we achieve a PCC of 0.86 and a RMSE of 1.19 kcal/mol, outperforming both iSEE and MuPIPR (Table 2). On S2003, we compared MuLAN results with those from DLA–Mutation (Mohseni Behbahani *et al*., 2023), a model that uses 3D convolutions and a self-supervised pre-training objective based on masking to model the local structural environment around the mutated residue, requiring high-resolution complexes for training and testing. In spite of a total different approach, based on global sequence features, we achieve a PCC of 0.79 in mutation-based cross validation, slightly lower than 0.81 of DLA–Mutation, and a PCC of 0.685 in the same complex-based train/test splitting considered. While being significantly lower than the 0.735 achieved by DLA–Mutation when enriched with auxiliary features (namely the structural region, evolutionary conservation, GEMME score, buriedness and physico-chemical properties of the wild-type residue), the result is comparable with the base model implementation (DLA–Mutation* in Table 2), that considers only the structural region as auxiliary feature, and outperforms the DLA-mutation ablated version that includes all additional features, but is not pre-trained with the masking objective (PCC = 0.608). These results show good generalization capabilities of the model, that do not depend on the accuracy and resolution of available structures in the region of interest. Finally, S1131 and S4169 are widely used evaluation datasets, constructed with single-point mutations from SKEMPI v1 and v2, respectively. On S1131, we outperform TopNetTree, Hom-ML-V2, and the more recent MpbPPI, all of which are based on deep learning, achieving a PCC of 0.87 and an RMSE of 1.20. However, on S4169, our model’s performance falls short, with a PCC of 0.739, which is significantly lower than the 0.79 obtained with TopNetTree and MpbPPI. As mentioned above, we partially attribute this loss in performance to the less informative embedding representation of some proteins in this dataset, such as antibody-antigen complexes.

#### PLMs fine-tuning improves performances for smaller models

In an attempt to enhance the model prediction capabilities, we experimented with fine-tuning of backbone pLMs and applied two alternative fine-tuning strategies, with different objectives:

- *evo-tuning* : this strategy involves continual learning on the masked language modeling (MLM) task, where the model predicts a masked fraction of residues in the sequence based on the context. We restricted the training dataset to the four most common species in SKEMPI, namely *Homo sapiens, Bos taurus, Mus musculus* and *Escherichia coli* ;
- *supervised fine-tuning* : this approach focuses on token classification fine-tuning for predicting residues that belong to inter-protein interaction surfaces. This information is not directly captured by the base models, despite their ability to predict intra-protein contact maps with minimal supervision (Rao *et al*., 2020).

We tested the fine-tuned models by using their embeddings as inputs to MuLAN, and we adopted a mutation-based cross-validation scheme on S1102 and S1400 datasets. The results for RMSE, reported in Table 3, demonstrate that the *evo-tuning* strategy can effectively lead to improved performances even for smaller models, like ESM2–35M, due to the higher taxonomic proximity between sequences appearing in the training and in downstream tasks. On the other hand, *supervised fine-tuning* on interface classification generally increases the RMSE and therefore is not beneficial for the ΔΔ*G* prediction.

**Table 3:** RMSE of MuLAN predictions for fine-tuned PLMs used as feature extractors. Model performances are evaluated on the benchmark datasets S1102 and S1400. “Baseline” refers to the pre-trained PLMs.

| PRE-TRAINED MODEL | BASELINE |  | EVO-TUNING |  | SUPERVISED FINE-TUNING |  |
| --- | --- | --- | --- | --- | --- | --- |
|  | S1102 | S1400 | S1102 | S1400 | S1102 | S1400 |
| ESM2-35M | 1.289 | 1.188 | 1.267 | 1.197 | 1.242 | 1.221 |
| ESM2-650M | 1.215 | 1.181 | 1.202 | 1.167 | 1.244 | 1.245 |
| ProtBert | 1.270 | 1.241 | - | - | 1.272 | 1.227 |

### Light attention weights focus on protein interaction sites

Protein interaction surfaces are plastic, adapting to different partners such that the same residue at the interface might contribute differently to the binding strength of each interaction. Therefore, evaluating which residues are more relevant in the context of specific partners is crucial for building a comprehensive understanding of these interactions. MuLAN contributes directly to this by providing insights into residue relevance and interaction strength based on the specific partners involved.

MuLAN learns a weighted average of residue representations through the application of convolutional filters, which are responsible for its light attention mechanism. This architectural design allows it to extract attention weights that offer physical insights into the interaction and provide a biological interpretation of the resulting predictions. The availability of experimental 3D structures in the SKEMPI database, combined with the per-residue processing of the model, enables a direct connection between the network’s attention outputs and the corresponding regions on the proteins. Namely, we conjectured that high-attention positions might be related to regions on the protein surface that are more likely to contribute to variations in binding affinity. In order to test this hypothesis, we computed the attention scores for S1102 dataset, for MuLAN–Ankh– large and MuLAN–ESM2–3B models, and compared them with interfaces extracted from the corresponding Protein Data Bank (PDB) (Burley *et al*., 2023) files with INTBuilder (Dequeker *et al*., 2017). For attention weights of size (*N*_heads_, *N*_hidden_, *L*) We applied an average pooling over the first two dimensions, obtaining a score for each position that was then rescaled in the range [0, 1]. We found that such scores proved to be effective in the discrimination between regions that belong or not to the interface, especially for MuLAN-Ankh-large model (see Figure S1).

To test the generalization of this unsupervised method, we resorted to ProteinNet dataset, with interface labels extracted using LEVELNET (Behbahani *et al*., 2023). These interface labels refer to the union of known interacting surfaces for each protein, rather than to interactions within a specific complex. After removing sequences present in SKEMPI, we selected those with lengths between 30 and 1024 amino acids and with a fraction of residues in interface regions between 0.1 and 0.5. We then clustered the remaining sequences at 30% sequence identity using MMseqs, reducing the dataset from 44,000 to just under 10,000 sequences. From this reduced set, we randomly selected a subset of 500 sequences as the external test set. For comparison, besides attention scores and PLMs embeddings, we also extracted pLDDT scores of test sequences for AlphaFold predicted structures available on UniProt, that where shown to be informative about possible interface regions (Seoane and Carbone, 2022). By removing sequences without an associated structure and with discrepancies in the primary structure, we restricted the test set to 308 sequences. The ROC curve for the test set is reported in Figure 2C. While embeddings extracted from PLMs do not contain any general information about the interface region, as shown by the random-like behaviour of the ROC curve, attention weights extracted from MuLAN appear to capture a signal that generalizes beyond the complexes and specific mutations present in the training set. This is particularly true for MuLAN–Ankh, that achieves an AUC of 0.62, higher than the AUC associated to pLDDT scores. This result proves that attention weights from MuLAN model can effectively discriminate interface regions, even if the model is not trained on this specific task.

**Figure 2:**
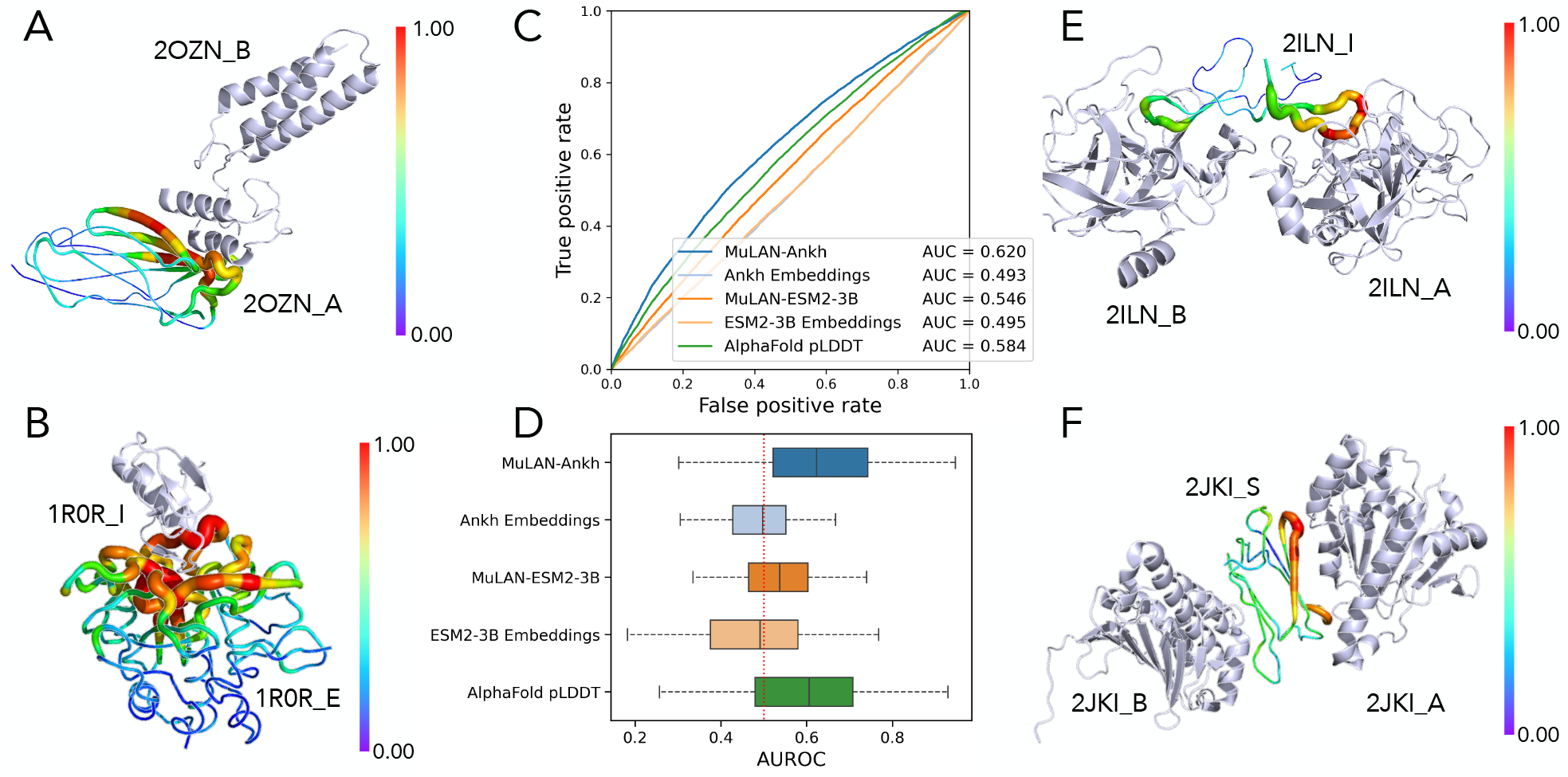
Predictions of interface residues from MuLAN learned representations. (A) Cohesin (2OZN_A) in complex with Dockerin (2OZN_B). Complex belonging to the ProteinNet test set. The color scale represents predicted MuLAN–Ankh probabilities for a residue to belong to the interface. (B) Predictions by MuLAN–Ankh on the Serine Protease (subtilisin Carlsberg, 1R0R E) and Kazal inhibitor (1R0R_I). Complex belonging to the S1102 dataset. Color scale as in A. (C) ROC curve for several models tested on 308 sequences randomly selected from the filtered ProteinNet dataset. The corresponding AUC is shown in the inset figure legend. (D) Distributions of AUROC values for the 308 proteins in the test set extracted from ProteinNet in C, for the different models considered. (E) Bowman-Birk inhibitor (2ILN_I) in complex with two bovine trypsin chains (2ILN_AB). Complex belonging to the ProteinNet dataset. Color scale as in A. (F) Heat shock proteins Hsp90 N-terminal (2JKI_AB) in complex with the adaptor protein Sgt1 (2JKI_S). Complex belonging to the ProteinNet dataset. Color scale as in A.

Figure 2AB illustrates MuLAN’s analysis on two binary complexes. For each protein, all residues are analyzed, with those displaying higher attention (in red) being the most relevant to the interaction. In Figure 2EF, the protein under analysis is in contact with two identical chains, with one interaction considered by MuLAN more relevant than the other, potentially suggesting a preference in the assembly order. Notably, in both cases, the interaction with the second chain is also identified (in green) and distinguished from non-interacting parts of the proteins (in blue). In all four examples, MuLAN focuses on the interaction surfaces of the complexes.

### MuLAN reconstructs complex-dependent mutational landscapes

The ability to rapidly predict ΔΔ*G* variations for any given mutation in a protein complex using only sequence inputs enables the reconstruction of Deep Mutational Scanning experiments that measure binding affinities in complex interactions. These reconstructed matrices not only assess the stability of the query protein upon mutation but are also informed by the interacting partner, resulting in key differences in predictions depending on the specific binary complex. For the first time ever, these mutational landscapes are computationally reconstructed as complex-dependent.

We explored the model capabilities in this scope by selecting some test cases and comparing the predictions with those obtained with ESCOTT (Tekpinar *et al*., 2024). In all the experiments, we used the MuLAN– Ankh–large model (thus without addition of *i*GEMME scores, to avoid possible biases), that was trained on S1102 dataset. To reduce information leakage, we ran BLASTp and removed complexes for which one of the partners shared more than 40% sequence identity with query proteins from the training dataset.

#### Residues involved in BPTI interactions identified as highly mutation-sensitive by MuLAN

We considered Bovine Pancreatic Trypsin Inhibitor (BPTI) interacting with three homologous proteins in binary complexes, namely its coevolved biological target bovine trypsin (BT), the noncognate trypsin paralogs bovine *α*-chymotrypsin (ChT) and human mesotrypsin (MT), taken from (Heyne *et al*., 2021). Sequences for the interacting proteins were taken from the corresponding PDB entries 3OTJ, 1CBW and 2R9P. BPTI protein presents a loop involved in the interaction, for which mutational effect are difficult to predict for tools based on single sequences.

We ran the trained model and obtained predictions for all possible single-point mutations in BPTI, and we ranksorted the predictions in order to compare them with ESCOTT, that assigns scores in the range [0, 1], with higher values related to more impactful mutations. Our findings are summarized in Figure 3, where we report MuLAN predictions as well as their comparison with ESCOTT. We also report a comparison with T_*JET*_ scores (Engelen *et al*., 2009) measuring the evolutionary conservation levels of the amino acids (Figure 3A). While the model fails at capturing the differences in experimental binding affinities for the three complexes due to high sequence similarity in interacting proteins, it is able to capture additional information on single amino acid variants predictions caused by interaction. In particular, MuLAN results tend to enhance scores for residues belonging to the interacting surface (see red residues and regions in Figure 3C, top), and decrease the peaks for other regions sensible to mutations see grey residues and regions in Figure 3C, top). The predictions also show general similarities with evolutionary conservation scores (Figure 3A), which is an information that is expected to be captured by PLMs (Meier *et al*., 2021). However, while a highly conserved region is found in the interior of the protein, and only a subset of residues involved in the interaction is labeled as conserved, MuLAN extrapolates beyond this conserved context, by assigning the highest scores to residues on the interface (in particular residues 34-38 and 11-15). This example, combined with the unsupervised analysis of interfaces from MuLAN representations, proves that the model can effectively learn information on the protein complex from single sequences.

**Figure 3:**
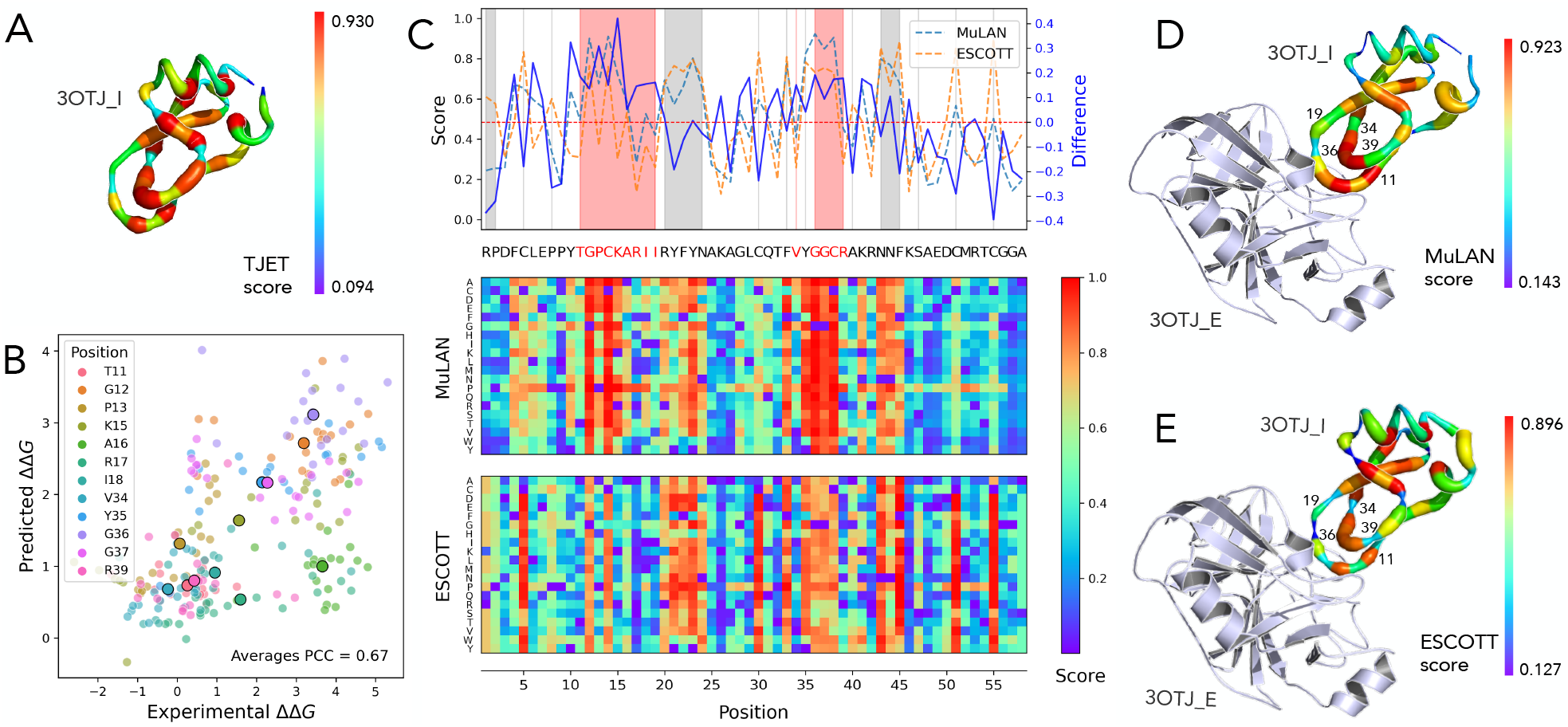
MuLAN analysis of the BPTI (Bovine Pancreatic Trypsin Inhibitor) complexed with trypsin. (A) T_*JET*_ evolutionary conservation scores for residues in BPTI, as computed by JET (Engelen *et al*., 2009) from the multiple sequence alignment, plotted on structure PDB ID 3OTJ I. (B) MuLAN ΔΔ*G* predictions for mutations considered in [(Heyne *et al*., 2021)], for which experimental values are available. The different colors refer to the positions of the mutated amino acid, and the highlighted dots represent the averages over all mutations in that specific position. (C) MuLAN and ESCOTT positional average scores (left axis) are plot together with their differences (right axis). Interface residues correspond to positions 11-19, 34 and 36-39 in the sequence and are indicated with red characters. They are highlighted either by a red line or a red background in the plot. They are identified from the complex structure PDB ID 3OTJ. Positions highlighted by a grey line or a grey background correspond to residues with average score, over all mutations, > .5 in ESCOTT, hence predicted to be highly sensitive to mutations. MuLAN (center) and ESCOTT (bottom) landscape reconstructions, where the color scale from 0 (low effect) to 1 (high effect) corresponds to MuLAN/ESCOTT scores. (D) Residues in the BPTI protein are colored with positional averages from the MuLAN prediction matrix in (C). (E) Residues in the BPTI protein are colored with positional averages from ESCOTT prediction matrix in (C).

#### MuLAN facilitates the selection of structural conformations in complexed Thioredoxin

The aim of this analysis was to determine if MuLAN could identify essential residues of Thioredoxin f2 (TRX-f2) from *Chlamydomonas reinhardtii*. Thioredoxins are widely spread across the tree of life and play a key role in controlling the redox status of protein disulfide bonds in all non-parasitic organisms (Collet and Messens, 2010). Thierodoxins exhibit a characteristic three dimensional structure classified as TRX fold, which is extremely conserved throughout evolution. Their activity is ensured by the presence of a solvent-exposed motif (most commonly Trp-Cys-Gly-Pro-Cys) containing two cysteine (Cys) residues that are able to catalyze protein disulfide reduction (Lemaire *et al*., 2018).

We examined TRX-f2 interactions with various partners in *C. reinhardtii*, including ferredoxin-thioredoxin reductase (FTR), sedoheptulose-bisphosphatase (SBPase), fructose-1,6-bisphosphatase (FBPase), and Phosphoglycerate kinase (PGK). These interactions are particularly challenging to study as the structures of these complexes remain undetermined. Similarly to the BPTI case, we predicted ΔΔ*G* values for all single-point mutations in TRX-f2 in complex with the considered partners (Figure 4A for the interaction with FTR). We compared the regions highlighted by this analysis Figure 4B) and by the attention mechanism, with the different interaction surfaces identified by AlphaFold–Multimer (Figure 4CD). We found that the model identified the same interacting surface of TRX-f2, not only according to the MuLAN–Ankh predictions that do not depend on the partner, but also according to the ΔΔ*G* for the different complexes. For the TRX-f2-SBPase complex, we generated the AlphaFold–Multimer and ESMFold complex structures (Figure 4D). Based on the mutational landscapes computed for both TRX-f2 and SBPase, MuLAN can easily indicate the most favorable complex, which in this case turns out to be predicted by ESMFold, due to the colocalisation of the interaction surfaces and the presence of necessary cysteines in the TRX-f2 binding region.

**Figure 4:**
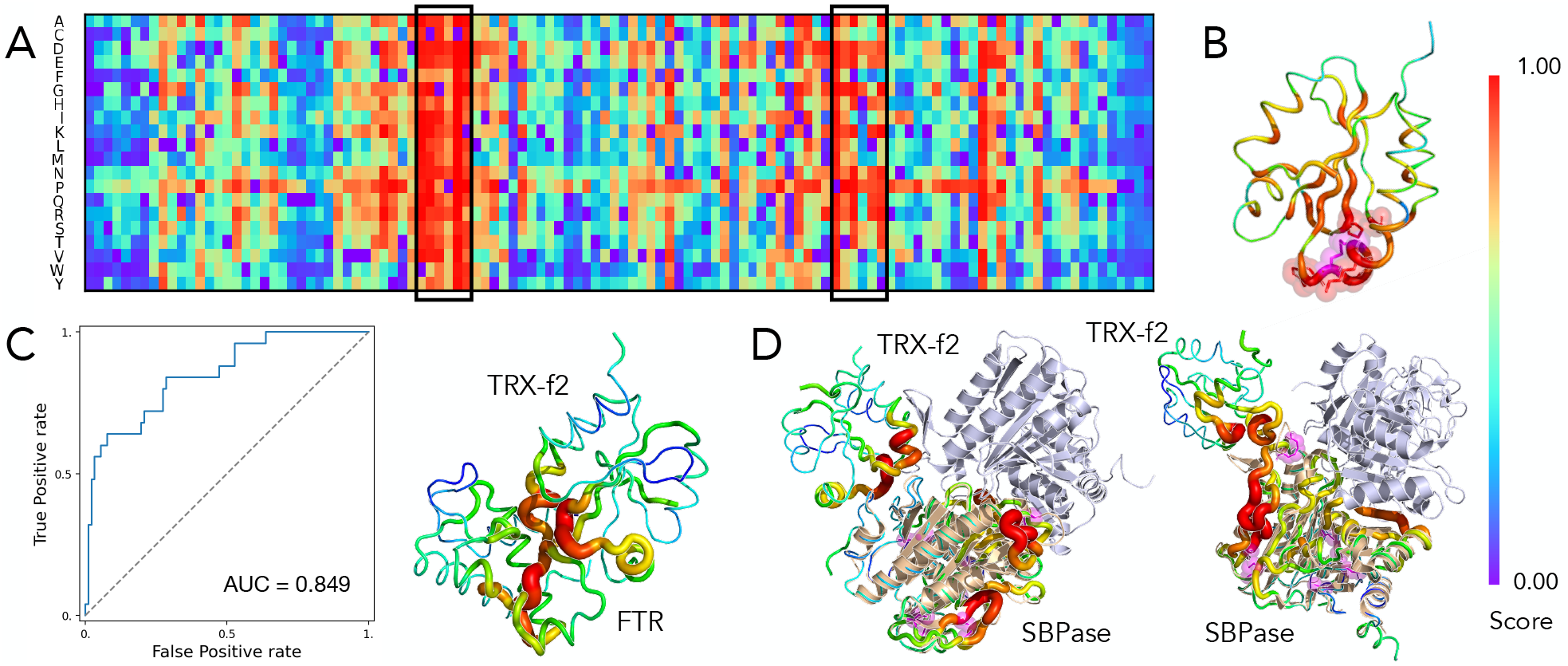
MuLAN predictions for TRX-f2 complexed with diversed partners. (A) Mutational landscape predicted by MuLAN for TRX-f2 in interaction with Ferredoxin-thioredoxin reductase (FTR), where highlighted regions are mostly involved in the interaction. (B) Average predicted values projected on TRX-f2 structure; the residues associated with highest values are represented as spheres, among which cysteins involved in redox catalyzation are highlighted in magenta. (C) Interface predictions for TRX-f2–FTR interaction, with associated ROC curve and projection on AlphaFold-Multimer predicted structure (RMSD=0.374 from homologous complex PDB ID 2PVO, in *Spinacia oleracea*). 27 residues in TRX-f2 were positively labeled as part of the interface, based on a 5.0 Å distance cutoff. (D) TRX-f2 complexed with SBPase, as predicted by AlphaFold-Multimer (on the left) and ESMFold (on the right), with projected interface predictions. SBPase in dimeric form (PDB ID 7B2O) is aligned to the predicted structure. Cysteines on SBPase, that suggest possible interaction sites, are highlighted in magenta.

#### MuLAN enhances the prediction of gain-of-function mutations in the MeFV gene

Gain-of-function mutations are a class of missense mutations leading to enhanced protein activity. Most of the tools for classification of mutations, including PRESCOTT and AlphaMissense, struggle with the identification of pathogenic gain-of-function mutations, in particular for those found in the MEFV gene, involved in various autoinflammatory diseases. This protein is of particular interest since many single-point gain-of-function mutations cause Familial Mediterranean Fever (FMF) and no current computational approach performs well on it. In an attempt to predict the effects of such mutations, we reconstructed the mutational landscape matrix with MuLAN for MEFV protein, by considering its homodimeric interaction (Figure 5BC). For comparison, we tested the same procedure also on other proteins from the GoF dataset known to be involved in autoinflammatory diseases. The dataset contains single-point mutations for 13 proteins, in variable proportions (from the single mutation R219H of CEBPE to the 30 mutations of IFIH1). We compared the results obtained with PRESCOTT, ESCOTT and AlphaMissense on this set of mutations, and extracted the attention scores from MuLAN for interface prediction (Figure 5A). MuLAN improves over all computational methods for MeFV and outperforms AlphaMissense on almost all other proteins in the dataset. Figure 5B illustrates MuLAN evaluation of a MeFV region where a number of known pathogenic mutations is localised and the corresponding ESCOTT evaluation which misses to highlight highly sensitive residues.

**Figure 5:**
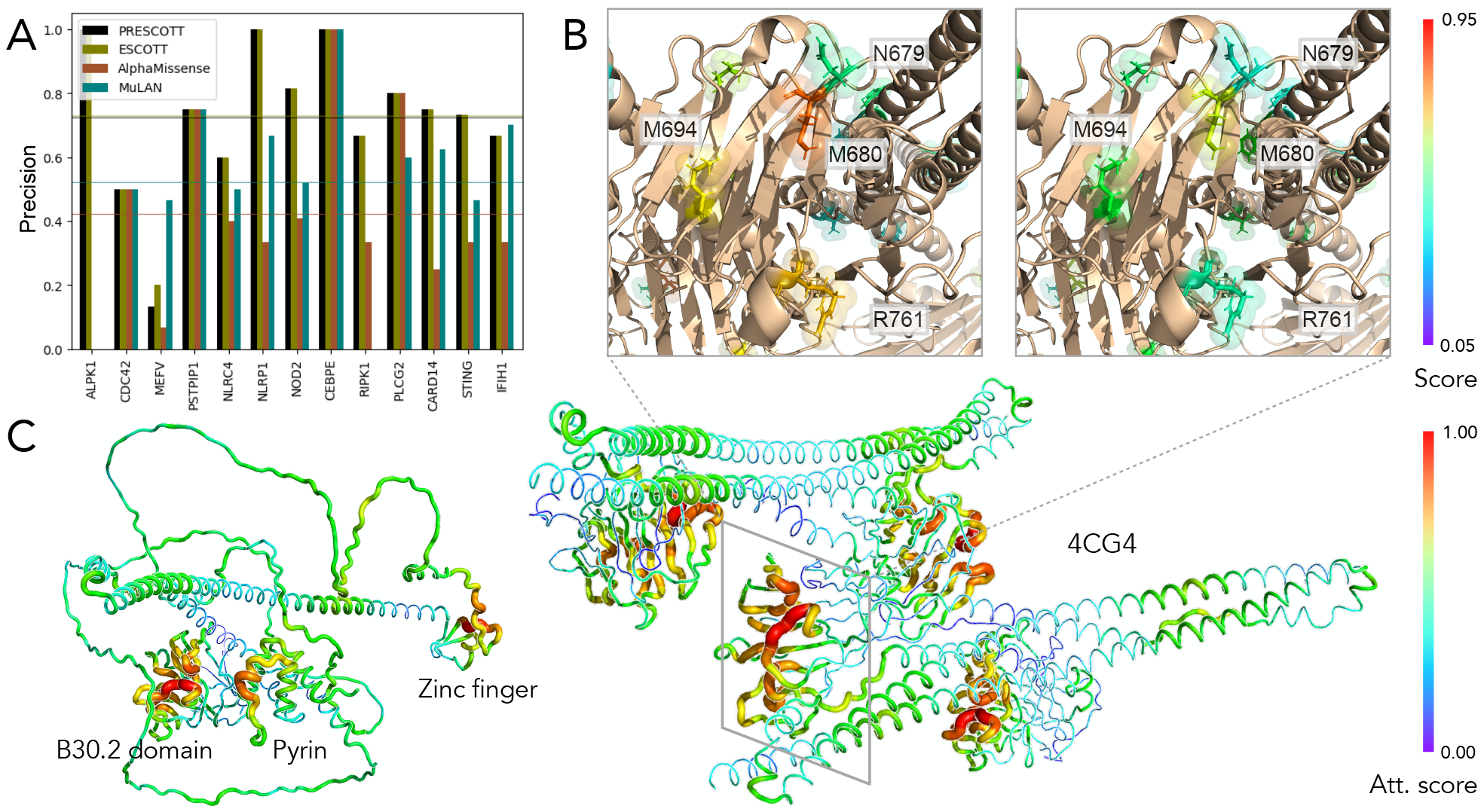
MuLAN predictions for Gain-of-Function (GoF) mutations in MEFV. (A) Fraction of correctly predicted pathogenic mutations from the GoF dataset (Tekpinar *et al*., 2024), comprising 13 proteins involved in inflammation. (B) Normalized average predictions from MuLAN (left) and ESCOTT (right) for some of the position considered for GoF mutations in the MEFV protein. (C) Interface scores computed from MuLAN and projected on AlphaFold predicted structure of the MEFV protein (left), where the different domains are highlighted, and on homodimeric complex assembly through the CHS-B30.2 domain (PDB ID 4CG4), corresponding to residues 414-781 of MEFV. MuLAN analysis of MeFV was performed on the dimeric complex.

### Computational details

Executing the MuLAN model requires neither accurate structures of the complexes nor the pre-extraction of specific features from the input, which could prove a bottleneck for structure-based approaches. On the contrary, the only required inputs are the sequences of the interacting protein complex, that are then tokenized and embedded in the PLM, and the specific mutations to evaluate. Once the embedding for mutated proteins are computed, the average evaluation time for the downstream model is around 0.015 second on a A100 GPU for one mutation, scaling linearly with the length of input sequences. The computation of embeddings is more time-consuming, leading to an average execution time of 1 second per mutation for the end-to-end model. However, for the reconstruction of entire mutational landscapes, time performances improve by pre-computing embeddings for the protein of interest in batches. In practical terms, for the reconstruction of entire mutational landscapes, 1,000 amino acids long proteins require about 1h while proteins of 200 amino acids roughly a minute.

## Methods

### Sequence embeddings

The primary structure of a protein of length *L* is a sequence of amino acids *S* = (*a*_1_, …, *a*_*L*_). In order to use it as input to a deep learning algorithm, it has to be encoded in a numerical format. The most unbiased and straightforward way to encode sequences without any prior knowledge injected is by representing each amino acid as a 1-hot vector, where the protein becomes a concatenation of *L* such vectors. In more recent years, the development of increasingly Large Language Models (LLMs) trained on protein sequences allows for contextualized embeddings. These protein Language Models (pLMs), often based on the transformer architecture (Vaswani *et al*., 2017), and capable of scaling up to billions of parameters, utilize a multi-head attention mechanism to capture long-distance correlations between amino acids. By predicting a fraction of masked residues from the context given by the rest of the sequence over millions of sequences and iterations, the model learns high dimensional representations (size *H*, typically of the order of 10^3^) for each amino acid. Their concatenation defines the protein embedding, of dimension *L × H*.

In many protein-level downstream tasks, such as solubility, localization or fitness prediction, an embedding that is independent of the length of the protein is required, which is usually obtained by global pooling over the length dimension (Rives *et al*., 2021). Dimensional reduction techniques applied to the extracted embeddings reveal that the models effectively learn some local features of single amino acids, for example hydrophobicity, charge, polarity, size, and global features of the whole proteins, such as conserved regions, location in the cell, and even structure and function to some extent (Lin *et al*., 2023; Rao et al., 2020). Similarly to LLMs, the performances of pLMs in zero-shot or few-shot downstream tasks are a measure of the information contained in the extracted embeddings and, ultimately, of the effectiveness of the encoder model itself. Up to date, pLMs have been developed in order to encode (encoders) or generate (decoders) single protein sequences, with performances that generally scale with the size of the model, but how to combine these representations in order to encode and predict properties for a protein complex still remains a challenge, and may likely depend on the considered task.

### A deep learning architecture with light attention

To extract and combine information from the embeddings of both wild-type and mutated proteins, we developed an architecture incorporating a Siamese encoder and a feature-combination module to perform the final regression. The model is structured into three parts: (i) a pre-trained protein language model with frozen weights, employed to extract contextualized embeddings from the input sequences; (ii) an attention block that processes the embeddings to generate vectors whose size does not depend on the length of the sequence; (iii) a fusion block designed to combine the encoded inputs for predicting the target property (Figure 6).

**Figure 6:**
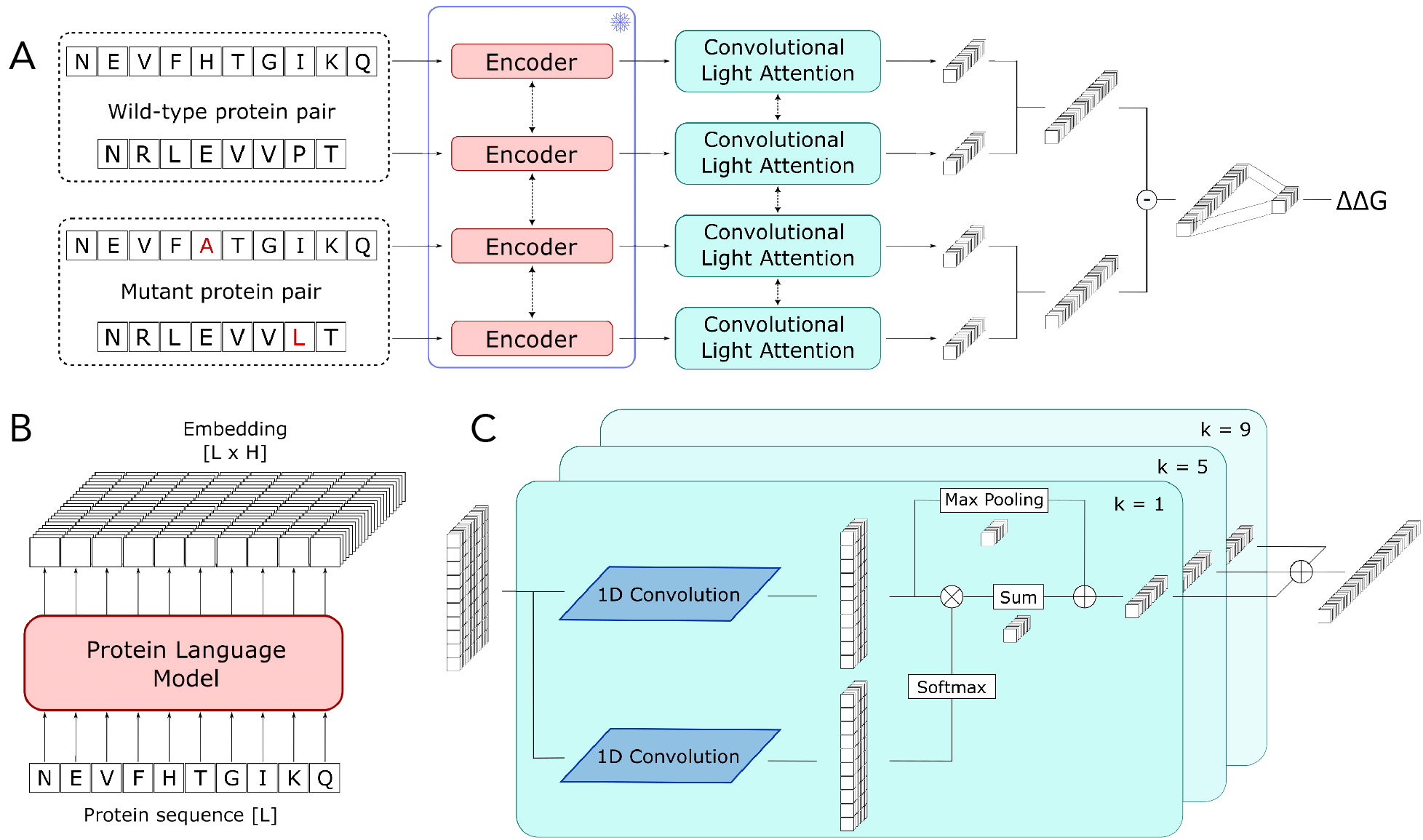
Representation of the Light Attention model architecture. (A) MuLAN full architecture: embeddings for both wild-type and mutated proteins are extracted from a pLM encoder, with frozen weights (B); each embedding undergoes processing within a Siamese Light Attention block, which normalizes weights across residues and learns a vector representation (C); these individual representations are subsequently combined pairwise to predict ΔΔ*G*.

The model processes the four input sequences independently, by applying a Light Attention Block (Stärk *et al*., 2021) on each. The architecture is built upon two parallel 1D convolutional layers applied to each sequence (Figure 6C). Of these layers, *C*_1_ and *C*_2_, one applies a dimensionality reduction of the input representation, cutting down the latent dimension (corresponding to the number of “channels” in convolution) from around 10^3^ to 64, and the other is followed by a SoftMax function applied across the length dimension, thus assigning a normalized weight to each residue within each latent dimension separately. We proceed to perform a point-wise product of the two outputs, summing over the length dimension, according to the equation

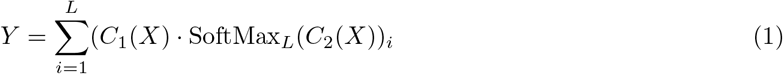

The result is a local linear attention mechanism, the scope of which can be varied by modifying the kernel size *k*. Testing different values for *k*, we discovered optimal performance with blocks applied in parallel for *k* = [1, 5, 9], and by joining their outputs together. Additionally, we applied a global MaxPooling over the length dimension for each convolutional head, which is then concatenated with the representation extracted directly from the Light Attention mechanism. The resulting flattened vector is then fed into a multilayer perceptron (MLP) layer equipped with a LeakyReLU activation function prior to entering the fusion block.

In the last layer, the outputs for the two interacting proteins are merged using both the dot product and the absolute value of the difference of the embedding vectors, in order to ensure that the resulting representation remains symmetrical regardless of the order in which the two proteins are considered. The final embeddings for both the wild-type and mutated complex are then subtracted from one another and fed into a linear layer to predict ΔΔ*G*. A schematic representation of the complete model is depicted in Figure 6.

The training goal is to reduce the mean squared error (MSE) between the observed experimental values and the model’s predictions. The evaluation of the models involves calculating the Pearson correlation coefficient (PCC), which assesses the linear correlation between the experimental outcomes and the predictions, and the root mean squared error (RMSE) defined as:

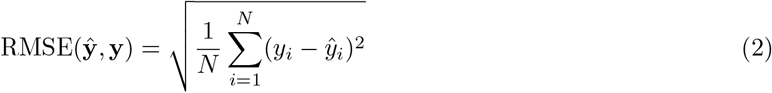

where **ŷ** represents the actual values and **y** denotes the predicted values.

### pLMs fine-tuning

To enrich the information content of the embeddings derived from pLMs, we designed two fine-tuning strategies using smaller-scale models: masked language modeling and binary classification at the single residue level. In the first approach, following (Zhou *et al*., 2020), we adopted the same self-supervised task that was used for pre-training the model. However, we narrowed down the training dataset to sequences exclusively from the most frequently occurring species in our dataset, specifically *Homo sapiens, Bos taurus, Mus musculus*, and *Escherichia coli*. This strategy aimed to tailor the model’s focus towards sequences more closely related to those found in the protein complexes under study. We named it “evo-tuning”.

The second fine-tuning strategy, named “supervised fine-tuning”, involves supervised binary classification, distinguishing whether residues are part of the protein’s interacting surface. To label the training data, we utilized LEVELNET software (Behbahani *et al*., 2023) on all the proteins in the dataset, excluding those listed in SKEMPI v2.0 (Jankauskait *et al*., 2018). LEVELNET identifies interface regions from online databases for each biological entity, and maps them to similar sequences through homology transfer. Hence, for a protein, labels are defined on the union of the interface regions known to LEVELNET. Out of nearly 77,000 sequences evaluated, about 28,000 were found to lack an interacting surface, possibly due to insufficient data, and were therefore excluded from the dataset. This process resulted in approximately 49,000 sequences, which were randomly divided into training and test datasets using an 80/20 split ratio. This approach is driven by the observation that mutations significantly impacting ΔΔ*G* often occur at the interface. Incorporating this information into the embeddings is expected to enhance the training efficiency of the downstream model.

### Extraction of Attention weights

The model’s design is crafted to promote interpretability by retrieving attention weights from its layers. Outputs from the attention heads are extracted and concatenated along the position dimension, such that each residues is encoded in a vector of dimension *N × C*, where *N* is the number of heads and *C* is the number of filters in the convolution (in our case, 3 and 64 respectively). By averaging over the obtained vectors, and applying MinMax normalization to the result, we allocate a singular scalar weight to each residue. This weight reflects the significance of that residue’s representation in determining the final ΔΔ*G* prediction.

### Datasets

#### ΔΔ*G* **experimental data**

The main source of our data is the SKEMPI v2.0 database, which is widely used in the context of the prediction of binding affinity changes upon mutations (Jankauskait *et al*., 2018). It is the most complete database for experimentally measured binding affinities for wild type and mutated complexes, for which the structure is solved and available in the Protein Data Bank (PDB) (Burley *et al*., 2023). SKEMPI v2.0 reports measurements for 7085 single-point and multiple mutations obtained from 345 different complexes, and it includes other smaller databases such as AB-Bind (Sirin *et al*., 2016), PROXiMATE (Jemimah *et al*., 2017), and dbMPIKT (Geng *et al*., 2019b). For each entry, the PDB structure of the wild-type complex is provided, as well as the names of the considered interacting partners in the complex, the mutated residue(s), the experimental methods employed for each measurement, and the associated affinities for wild-type and mutated complex, *k*_*wt*_ and *k*_*mut*_. From them, Δ*G* can be computed as

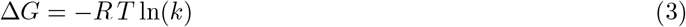

where *R* is the molar gas constant and *T* is the absolute temperature.

The structural region of the mutation site is also reported, as belonging to one of the five classes COR, SUP, RIM (corresponding to core, support and rim regions at the interface), INT or SUR (for interior or surface) (Levy, 2010). The tendency is towards higher variations in binding affinity for mutations located in the interacting surface, where most of them tend to be deleterious rather than beneficial.

The intrinsic asymmetry of the database must be considered during the model development. This can be addressed either by deliberatly overlooking the asymmetry, resulting in a predisposition for positive predictions, or by augmenting the database with inverse mutations, as in (Wang *et al*., 2020), coupled with incorporating a distinct term in the loss function to attain symmetry. Although this method can enhance performance in cross-validation assessments, it may lead to higher model redundancy, thereby reducing its ability to generalize to unseen data.

The SKEMPI v1.0 and v2.0 databases have been a source for numerous benchmark datasets used in recent years to train and evaluate a variety of models grounded in physics, statistics, and more recently, machine learning and deep learning approaches. We have also utilized some of these datasets for our training and evaluation, including S1400 featuring both single-point and multiple-point mutations, and S1102, S1131, S2003, and S4169 featuring only single mutations. The dataset names’ numbers reflect the count of data points, with each representing a complex associated with a ΔΔ*G* value linked to one or several mutations in one or both sequences. Table S1 reports the number of complexes, the number of mutations and the source of each dataset.

#### Enhancing pLMs with targeted fine-tuning

For the fine-tuning of pLMs, we leveraged datasets from two distinct sources. The dataset for masked language-modeling fine-tuning was sourced from STRING database (Szklarczyk *et al*., 2023), which reports over 20 billion experimentally measured and inferred protein-protein interactions across approximately 12,000 different organisms, amounting to more than 59 million protein sequences. We selected only proteins belonging to four species (*Homo sapiens, Bos taurus, Mus musculus* and *Escherichia coli*), resulting in a total of 66235 sequences for training and validation.

For the purpose of interface classification, we utilised the ProteinNet12 dataset, which offers a comprehensive collection of sequences, structures and multiple sequence alignments (AlQuraishi, 2019). We ran LEVELNET on more than 77,000 sequences clustered at 90% sequence identity from ProteinNet12, selecting a 90% sequence identity threshold for homology transfer. By removing sequences that were not present in LEVELNET database or for which no interface was found, we obtained interface labels for 48,452 proteins. The vast majority of these sequences have lengths shorter than 1024 residues, and longer sequences were randomly cut to this limit for training. A small set of 500 proteins from this reduced dataset was randomly selected for testing the prediction of interfaces from MuLAN attention.

### Implementation details

All the models, training and evaluation scripts are written in Python using the PyTorch library. Pretrained PLMs are available in the HuggingFace Hub and can be easily downloaded following the related documentation.

For efficient model training, we pre-computed the embeddings from pLMs for every wild-type and mutated protein in the training dataset. The single embeddings were then loaded in batches and dynamically zero-padded to the longest sequence length in the batch. iGEMME or log-odds scores were also pre-computed and stored separately, and then loaded and passed to the model with different modalities depending on the strategy adopted (single scores, averaged scores, etc.).

All downstream models are trained until convergence, using AdamW optimizer with default parameters, a batch size of 32 and an initial learning rate of 5 × 10^−4^, that is reduced by half on validation loss plateaux (with a patience of 5 epochs). The random seeds for dataset splitting in cross-validation experiments and the initialization of model parameters were set to 42 for reproducibility.

For pLMs fine-tuning, instead, we ran the training for 3 epochs with data parallelism across 4 A100 GPUs and gradient accumulation, with a total batch size of 256 and a learning rate of 5 × 10^−5^, with AdamW optimizer. Training data for fine-tuning tasks were randomly split in training and validation set, following a 0.8 - 0.2 ratio.

### Software availability

MuLAN is available at http://gitlab.lcqb.upmc.fr/lombardi/mulan under the [CC-BY-NC-SA 4.0 License](https://creativecommons.org/licenses/by-nc-sa/4.0/legalcode.en).

## Discussion

Understanding the physical interactions between proteins and other molecules is paramount for unraveling the molecular mechanisms that explain phenotypes. Our study focuses on protein-protein interactions leveraging deep learning to analyze sequence-derived protein representations, offering a robust alternative to structure-based methods. Deep learning enables us to tackle essential questions such as reconstructing interaction networks, identifying binding interfaces, and estimating changes in binding affinity. The sequence-based approach is especially advantageous for studying disordered and flexible regions involved in transient interactions, which are challenging to model with structural data. By extracting information from sequences, deep learning leverages evolutionary constraints that shape protein structure and function. MuLAN makes significant strides in this area, in three main ways: 1. it estimates the strength of interaction with binding partners, crucial for understanding disease susceptibility and drug efficacy. 2. it reconstructs a deep mutational scanning matrix for each interacting partner, incorporating information from both proteins. 3. it enhances the identification of residues in sequences that are involved in protein interaction sites, addressing the complex problem of varying interaction sites with different partners.

### MuLAN brings protein interactions into sharp focus

By analyzing a pair of interacting protein sequences, and by coupling it with the same pair “perturbed” with mutations at one or more amino acids, MuLAN can estimate changes in binding affinity. It does it at large-scale since it does not require precise complex structures. This approach enables the reconstruction of mutational effects on each protein and identify interaction sites, providing deeper insights into protein function and interactions. Notably, the mutational landscape of a protein varies when analyzed with different partners, demonstrating MuLAN’s ability to capture distinct interaction details for each protein pair. This capability closely reflects biological contexts, enhancing predictive accuracy beyond existing computational methods.

The significance of MuLAN is twofold. Firstly, it enhances the identification of residues in sequences that are part of protein interaction sites, offering an accurate method to address a major challenge in the field. Secondly, and most importantly, MuLAN introduces an innovative approach to reconstructing a deep mutational scanning matrix that considers protein interactions and incorporates information from both interacting partners. This hypothesis, previously unexplored, brings us closer to accurately modeling biological contexts. Unlike existing computational methods that overlook specific partner interactions, limiting their predictive capabilities, our approach identifies key differences by incorporating these interactions, thereby enhancing predictive accuracy and advancing our understanding of protein dynamics.

In summary, our study represents a significant advancement in the field of protein interaction research. By combining deep learning with sequence-based analysis, we provide a more comprehensive and accurate understanding of protein interactions and their underlying mechanisms. This not only enhances our ability to predict binding affinity changes but also opens new avenues for studying the complex nature of protein interactions in biological systems.

### Challenges in ΔΔ*G* prediction accuracy

As it is usually the case for machine learning regression models, extrapolation is challenging, as it requires the model to generalize well beyond the training data, and it is particularly true in the case where the training set is limited. In our endeavor to predict ΔΔ*G*, the training dataset predominantly feature wildtype and mutant complexes with binding differences under 10 kcal/mol (absolute value), with most data points below 4 kcal/mol and a handful of outliers. Additionally, there is a notable experimental preference for data on deleterious mutations over beneficial ones, leading to data asymmetry. These two factors contribute to the model’s inclination to predict lower positive values for unseen data, potentially misrepresenting the true ΔΔ*G* values. Nonetheless, we anticipate accurate ranking of predictions, where higher values signify deleterious mutations, and lower values indicate neutral or beneficial mutations. The pooling with other orthogonal methods, as we did here with the addition of iGEMME scores or, as recently shown (Lu *et al*., 2024) with AlphaFold3 ranking scores (Abramson *et al*., 2024), may help mitigating this limitation.

## Acknowledgments

Sorbonne Center for Artificial Intelligence (SCAI) for a PhD fellowship (GL); the Institut Universitaire de France (AC). This work was performed using HPC resources from GENCI–IDRIS (Grant 2022-102743) and Sorbonne University GPU clusters from SCAI and LIP6 laboratory (UMR 7606, Sorbonne University-CNRS).

## Supplementary Information

**Table S1:**
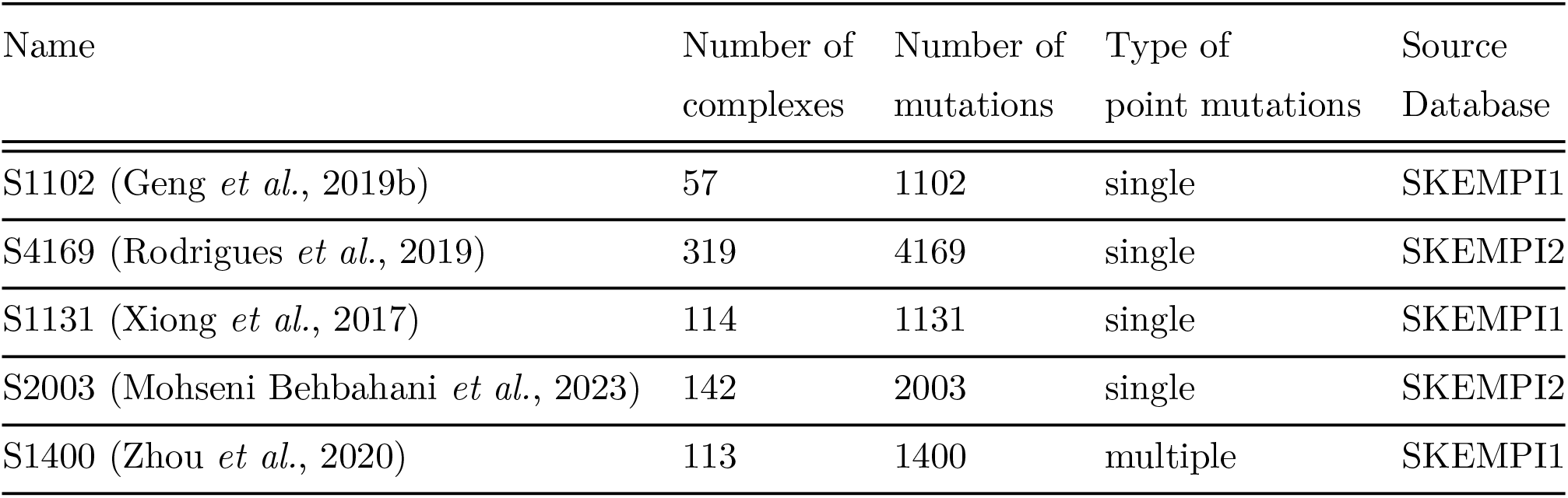
Benchmark datasets of changes of binding affinity upon mutation.

| Name | Number of<br>complexes | Number of<br>mutations | Type of<br>point mutations | Source<br>Database |
| --- | --- | --- | --- | --- |
| S1102 (Geng *et al.*, 2019b) | 57 | 1102 | single | SKEMPI1 |
| S4169 (Rodrigues *et al.*, 2019) | 319 | 4169 | single | SKEMPI2 |
| S1131 (Xiong *et al.*, 2017) | 114 | 1131 | single | SKEMPI1 |
| S2003 (Mohseni Behbahani *et al.*, 2023) | 142 | 2003 | single | SKEMPI2 |
| S1400 (Zhou *et al.*, 2020) | 113 | 1400 | multiple | SKEMPI1 |

**Figure S1:**
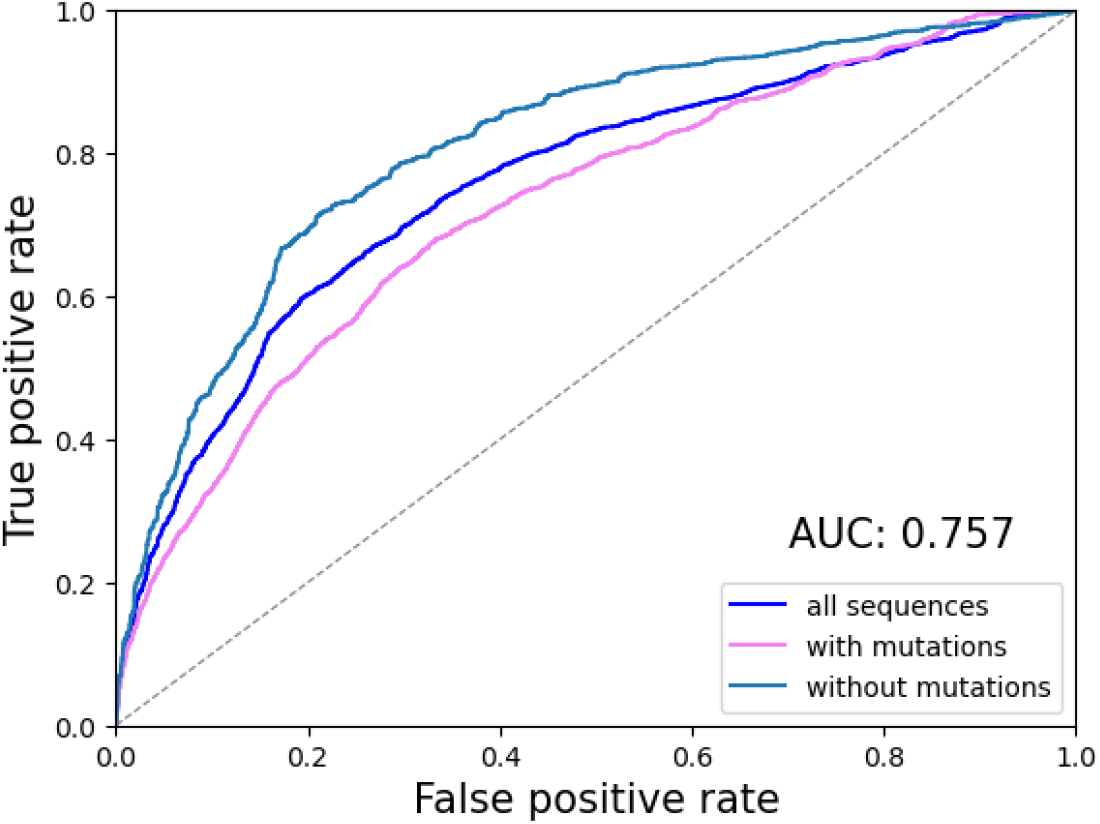
ROC curve for MuLAN-Ankh-large attention weights over S1102 training set. Attention weigths achieve an average AUC of 0.757 for all sequences, that does not appear to be biased on regions in the dataset where mutations are found (AUC = 0.809 for sequences without mutations, AUC = 0.723 for sequences that present at least a mutation).

